# Meiotic crossover patterning in the absence of ATR: Loss of interference and assurance but not the centromere effect

**DOI:** 10.1101/143651

**Authors:** Morgan M. Brady, Susan McMahan, Jeff Sekelsky

## Abstract

Meiotic crossovers must be properly patterned to ensure accurate disjunction of homologous chromosomes during meiosis I. Disruption of the spatial distribution of crossovers can lead to nondisjunction, aneuploidy, gamete dysfunction, miscarriage, or birth defects. One of the earliest identified genes in involved proper crossover patterning is textitDrosophila mei-41, which encodes the ortholog of the checkpoint kinase ATR. Analysis of hypomorphic mutants suggested the existence of crossover patterning defects, but it was not possible to assess this in null mutants because of maternal-effect embryonic lethality. To overcome this lethality, we constructed *mei-41* null mutants in which we expressed wild-type Mei-41 in the germline after completion of meiotic recombination, allowing progeny to survive. We find that crossovers are decreased to about one third of wild-type levels, but the reduction is not uniform, being less severe in the proximal regions of *2L* than in medial or distal *2L* or on the *X* chromosome. None of the crossovers formed in the absence of Mei-41 require Mei-9, the presumptive meiotic resolvase, suggesting that Mei-41 functions everywhere, despite the differential effects on crossover frequency. Interference appears to be significantly reduced or absent in *mei-41* mutants, but the reduction in crossover density in centromere-proximal regions is largely intact. We propose that crossover patterning is achieved in a stepwise manner, with the crossover suppression related to proximity to the centromere occurring prior to and independently of crossover designation and enforcement of interference. In this model, Mei-41 has an essential after the centromere effect is established but before crossover designation and interference occur.

## Introduction

Meiotic crossovers are subject to numerous mechanisms of spatial control to ensure proper disjunction of homologous chromosomes and generation of genetic diversity. Sturtevant (1913) described the phenomenon of crossover interference, where the presence of one crossover reduces the probability of crossovers nearby (reviewed in Berchowitz and Copenhaver 2010). Mather (1937) pointed out that for small chromosomes “the chiasma frequency equals one, no matter what the size;” Owen (1949) referred to this as the “obligate chiasma.” The phenomenon in which every pair of homologous chromosomes has at least one crossover that generates a chiasma to promote disjunction is commonly called crossover assurance (reviewed in Wang *et al.* 2015). Together with crossover homeostasis, which buffers crossover formation from increases or decreases in potential crossover precursors (Martini *et al.* 2006), assurance and interference demarcate the minimum and maximum number of crossovers per meiosis. Modeling suggests that crossover assurance, interference, and homeostasis are the result of a single patterning process (Wang *et al.* 2015). The mechanisms that achieve assurance, interference, and homeostasis remain obscure.

Less attention has been paid to the centromere effect, a spatial crossover patterning phenomenon first described by Beadle (1932). Crossovers are excluded from the vicinity of centromeres in many organisms, presumably because very proximal crossovers can interfere with homolog disjunction (*e.g.,* Koehler *et al.* 1996; Lamb *et al.* 1996). There are two components to the reduction in crossovers near the centromere. First, Muller and Painter (1932) reported that crossing over is absent or extremely rare within the “inert regions”, now known to comprise heterochromatic pericentromeric satellite sequence. The second component, which we refer to as the centromere effect, is the phenomenon Beadle described: the reduction in crossing over within crossover-competent regions of the genome as a function of proximity to the centromere. Beadle noticed that when regions with high crossover density were moved closer to the centromere by chromosome rearrangement, crossover frequency decreased. The converse – increased crossover density when centromere-proximal regions are moved away from the centromere – was shown by Mather (1939).

The mechanisms underlying the centromere effect are also unknown. Meiotic recombination is initiated by formation of DNA double-strand breaks (DSBs). Each DSB can be repaired into a crossover or a non-crossover; the latter can be detected when they result in gene conversion, the unidirectional transfer of sequence from a donor (a homologous chromosome) to a recipient (the chromatid that received the DSB). In *Drosophila*, DSBs appear be to be excluded from the pericentric heterochromatin, explaining the absence of crossovers on those regions (Mehrotra and McKim 2006). The centromere effect could in principle be explained by decreased DSB density in proximal regions. However, recent whole-genome sequencing reveals that the density of non-crossover gene conversion is relatively constant across the assembled genome on each arm (Comeron *et al.* 2012; Miller *et al.* 2016). This suggests that DSB density is also fairly constant across the chromosome arm and that the centromere effect is exerted by regulating the outcome of DSB repair (crossover or non-crossover) in a manner that is dependent on distance to the centromere.

Mutations in the *Drosophila mei-41* gene were first described by Baker and Carpenter (1972), who reported a polar reduction in crossovers, with a less severe effect on crossovers in proximal regions, and a possible decrease in interference. These observations suggest a potential role for Mei-41 in crossover patterning. Mei-41 is the *Drosophila* ortholog of ATR kinase, best known for regulating DNA damage-dependent cell cycle checkpoints (Hari *et al.* 1995). Consistent with this role, Mei-41 establishes a checkpoint that monitors progression of meiotic recombination (Ghabrial and Schüpbach 1999; Abdu *et al.* 2002). In addition, Mei-41 acts redundantly with ATM kinase to promote phosphorylation of histone H2AV at sites of meiotic DSBs (Joyce *et al.* 2011). However, it is unlikely that either of these functions explains the effects of crossover number or position noted by Baker and Carpenter. Understanding this role is further complicated by the finding that Mei-41 has an essential function in slowing the rapid nuclear cycles at the midblastula transition in embryonic development (Sibon *et al.* 1999). Females with null mutations in *mei-41* are sterile because this function is lost, and thus the alleles used in previous studies of meiotic recombination are either hypomorphic or separation-of-function (Laurençon *et al.* 2003).

We sought to investigate the possible function for Mei-41 in crossover patterning by analyzing crossover distribution in *mei-41* null mutants. To overcome the requirement for maternal Mei 41, we used a transgene in which *mei-41* expression is under control of a promoter that turns on only after recombination has been completed, thereby generating a fertile *mei-41* “meiotic recombination null” mutant. We find that crossover and non-disjunction phenotypes are more severe in this mutant than in previously-reported hypomorphic mutants. We observe a polar effect on *2L* but not on the *X*; we suggest that this is due to retention of the centromere effect, which is weak on the *X*. However, interference and assurance are greatly decreased or lost. We propose that the centromere effect is established early in the meiotic recombination pathway and that Mei-41 has a recombination role after this establishment but before interference and assurance are achieved. Loss of Mei-41 leads to exit from the meiotic recombination pathway after establishment of the centromere effect and repair is then completed by alternative mechanisms that lack interference and assurance. These findings provide insight into the establishment of crossover patterning.

## Materials and Methods

### Drosophila stocks

Flies were maintained at 25 C on standard medium. To overcome the maternal-effect embryonic lethality of *mei-41^29D^* null mutation (Laurençon *et al.* 2003; Sibon *et al.* 1999), wild-type genomic *mei-41* was cloned into the *P*{*attB, UASp, w^+m^*} vector. This construct was inserted into the genome via phiC31 integrase-mediated transgenesis, into the *X* chromosome landing site *M*{*3xP3-RFP.attP’*}ZH-2A (BestGene Inc., Chino Hills, CA). The resulting integrants, abbreviated herein as *M*{*UASp::mei-41*}, were crossed into a *P*{*mata4::GAL4-VP16*} background. All *mei-41* null assays used the genotype:

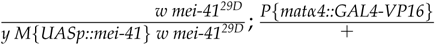

The *mei-41 mei-P22* double mutant was as above except the *3*^*rd*^ chromosomes were *mei-P22^103^st*/*mei-P22^103^Blm^D2^Sb P*{*mata4::GAL4-VP16*}. The *mei-9 mei-41* double mutant was as above except the *X* chromosomes were *y mei-9^a^mei-41^29D^*/*y M*{*UASp:: mei-41} w mei-9^a^mei-41^29D^*. The presence of the *mei-9*^*a*^ mutation was confirmed by allele-specific PCR and in genetic tests (see Table S5).

### Hatch rates

To test *M*{*UASp::mei-41*} rescue efficiency, 60 virgin females of appropriate genotypes were crossed to 20 isogenized Oregon-R^m^males (courtesy of Scott Hawley). Adults were mated in grape-juice agar cages containing yeast paste for two days prior to collection. Embryos were collected on grape-juice agar plates for five hours and scored for hatching 48 hours later.

### Crossover assays and analyses

Meiotic crossovers on *2L* were quantified by crossing *net dpp^d-ho^ dp b pr cn*/+ virgin females of the appropriate mutant background to *net dpp^d-ho^dp b pr cn* males. All six markers were scored in progeny from each genotype, with the of exception *mei-41*; *mei-P22*. In that case, 731 *XX* females were scored for all six markers and an additional 1023 *XXY* females and *XY* males were scored for *net—b*; eye color markers *pr* and *cn* were excluded because of the presence of a *w* mutation in the mothers. These data were pooled for a final *n* of 1754 progeny scored.

Meiotic crossovers on *X* were quantified by crossing *y sc cv v g f* · *y^+^*/*y* virgin females of the appropriate background to *y sc cv v g f* males. "· *y^+^*" is *Dp(1;1)sc^V1^*, a duplication of the left end of the *X*, carrying *y^+^*, onto *XR*. All six markers were scored in all progeny.

To measure chromosome *4* crossovers, the *mei-41* rescue genotype given above was made heterozygous for *PBac*{*y^+^w^+m^*}(101F) and *sv^spa-pol^*, which are near opposite ends of the assembled region of *4*. These females were crossed *w^1118^*; *sv^spa-pol^* males and the progeny were scored for the poliert eye phenotype associated with *sv^spa-pol^* homozygosity and the *w^+m^* of the *PBac* transgene. Although both the *M*{*UASp::mei-41*} and *P*{*mata4::GAL4-VP16*} transgenes also carry a *w^+m^*, both confer only mild eye coloration, so the strong red-eye phenotype of *PBac*{*y^+^w^+m^*}(101F) is easily discerned.

Genetic distances, expressed here in centiMorgans (cM) rather than “map units”, as is traditionally used in *Drosophila*, were calculated using the equations of Stevens (1936). Crossover density was calculated by dividing *cM* by the distance between markers (rounded to nearest 10 kb), using *Drosophila melanogaster* reference genome release 6.12 with transposable elements excluded, as described in Hatkevich *et al.* (2017). Including transposable elements in distances did not change any conclusions (see Tables S4a and S4b).

The coefficient of coincidence (*c*) is 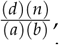 where *a* and *b* are the number of single-crossover progeny in two intervals being compared, *d* is the number of double-crossover (DCO) progeny, and n is the total progeny scored. This is equivalent to observed DCOs divided by expected DCOs if the two intervals are independent (no interference). Interference (*I*) is 1-*c*. Thus, *I* = 0 in the absence of interference and *I* = 1 if there is complete positive interference (no DCOs observed).

The centromere effect was quantified as in Hatkevich *et al.* (2017). The definition parallels that of *I*: 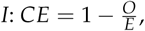, where *O* is the number of crossovers observed and *E* is the number expected based on the average crossover density across the entire region assayed. *CE* therefore describes the deviation in crossover density in in any interval from the mean density across all intervals.

For crossover assurance, we obtained the expected number of meiosis in which a give region of the genome (*X* or *net—cn*) had no crossovers (E_0_) from the Poisson distribution, using mean number of crossovers in that region as the average rate of success. To convert observed crossover classes (parental, single, double, and triple crossover) to bivalent exchange classes (E_0_, E_1_, E_2_, E_3_) we used the method of Weinstein 1936. This method accounts for the fact that an E_1_ “tetrad” gives two crossover chromatids and two parental chromatids, so the probability of recovering the crossover in the progeny is 0.5. Weinstein tested models with and without sister chromatid exchange and with and without chromatid interference (*i.e.*, whether the chromatids involved in the two crossovers of a DCO are independent of one another). We used the model that he found to be the best fit to two large *Drosophila* datasets: no sister chromatid exchange and no chromatid interference.

*X* nondisjunction was scored by crossing virgin mutant females of the appropriate genotypes to *y sc cv v g f* /*Dp(1:Y)B*^*S*^ males. Exceptional progeny for *X* nondisjunction events originate from diplo-*X* and nullo-*X* ova, resulting in *XXY* (Bar-eyed females) and *XO* (wild-eyed males) progeny, respectively. Numbers of exceptional progeny were doubled to account for *X* those that do not survive to adulthood (*XXY* and *YO*).

### Statistical analyses

For *cM* and *I*, 95% confidence intervals were calculated as 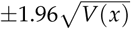, where *V(x)* is the variance of parameter *x*. 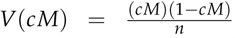 and 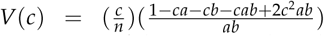 (Stevens 1936). For nondisjunction, 95% confidence intervals and comparisons of rates across genotypes followed the statistical methods developed by Zeng *et al.* (2010).

For between-genotype comparisons of interference we conducted *χ*^2^ tests on 2-by-2 contingency tables of observed and expected DCOs for each genotype. A 2-by-2 table is appropriate for counts of events that are positive integer values and for which there is an expectation under the null hypothesis that mutant and wild type have the same levels of interference given their levels of recombination. This expected number of DCOs is derived by applying a model of the frequency of double crossovers under no interference. Since the data do not have covariates or repeated measures, a *χ*^2^ test is the most straightforward. We applied Yates’ continuity correction because of low counts in some categories. A similar argument holds for the centromere effect and assurance. For assurance, we compared observed and expected E_0_ and E_>0_ classes. *χ*^2^ tests were conducted using the GraphPad QuickCalcs online tool (https://www.graphpad.com/quickcalcs/contingency1.cfm).

### Data Availability

The authors state that all data necessary for confirming the conclusions presented in the article are represented fully within the article. *Drosophila* stocks are available upon request.

## Results

### Post-germarium expression of mei-41 rescues embryonic lethality and creates a meiotic recombination null

*Drosophila* females homozygous for null mutations in *mei-41* produce no viable progeny due to a requirement for maternally-deposited Mei-41 at the midblastula transition (Sibon *et al.* 1999). *Blm* null mutants also exhibit maternal-effect embryonic lethality (McVey *et al.* 2007). To study meiotic recombination in *Blm* null mutants, Kohl *et al.* (2012) expressed wild-type Blm under indirect control of the *alpha tubulin 67C* (*matα*) promoter via the Gal4>*UASp* system. This promoter is specific to the female germline, with expression initiating in the early vitellarium (Sanghavi *et al.* 2013), by which time recombination should be compete. In support of this expectation, crossover and nondisjunction assays on the occasional surviving progeny of *Blm* mutant females give similar results to those from embryos rescued by expressing *UASp::Blm* with the *matα4::GAL4-VP16* driver in *Blm* null mothers (McVey *et al.* 2007; Kohl *et al.* 2012; Hatkevich *et al.* 2017).

We used the same system to overcome the maternal-effect inviability of embryos from *mei-41^29D^* homozygous null females (see Materials and Methods). To quantify the extent of maternal *M{UASp::mei-41*} rescue, we compared hatch rates of embryos from wild-type, *mei-41^29D^*, and *M*{*UASp::mei-41*} *mei-41^29D^* with and without *P*{*matα4::GAL4-VP16*} (Table 1). Embryos from females homozygous for *mei-41^29D^* with or without *P*{*UASp::mei-41*} but lacking *P*{*matα4::GAL4-VP16*} did not survive to hatching, whereas embryos from females with both components of the Gal4>*UASp* rescue system had a hatch rate of 52.8%. Most or all of the residual lethality is likely due to aneuploidy resulting from high nondisjunction in *mei-41* mutants (13.6% *X* nondisjunction among progeny surviving to adulthood; Table S5). Larvae that did hatch survived to adulthood, allowing for analysis of the crossover patterning landscape in a *mei-41* null mutant. For simplicity, flies carrying this transgene system are denoted below as *mei-41^29D^* or *mei-41* null mutants.

**Table 1.** Hatch rates for embryos from *mei-41* mutants.

| Maternal Genotype | Hatched (%) | Total (n) |
| --- | --- | --- |
| wild type | 73.1 <sup>a</sup> | 2035 |
| *mei-41*<sup>29D</sup> | 0 | 527 |
| *M{UASp::mei-41} mei-41*<sup>29D</sup> | 0 | 837 |
| *M{UASp::mei-41} mei-41*<sup>29D</sup>; *P{matα4::GAL4-VP16}* / + | 52.8 <sup>b</sup> | 1187 |

<sup>a</sup> This number is lower than expected for wild type. The cause of this is unknown.
<sup>b</sup> The apparent lack of complete rescue may largely be the result of a high frequency of aneuploidy resulting from the absence of *mei-41* during meiotic recombination.

### Crossover reduction in *mei-41* null mutants

*Drosophila mei-41* was initially characterized as a meiotic mutant by Baker and Carpenter (1972). Hypomorphic *mei-41* alleles resulted in an overall 46% decrease in crossovers relative to wild-type controls, measured in five adjacent intervals spanning the entirety of *2L* and proximal *2R* (about 20% of the euchromatic genome). We measured crossovers in this same region in *mei-41* null mutant females and found a significantly more severe reduction of 67% (*p* < 0.0001; Figure 1A and 1C). Given the many functions of Mei-41 in mitotically proliferating cells, we wanted to determine whether the remaining crossovers were meiotic or possibly resulted from DNA damage within the pre-meiotic germline. As Mei-P22 is required to generate meiotic DSBs (Liu *et al.* 2002; Robert *et al.* 2016), any crossovers that are independent of Mei-P22 most likely result from damage occurring in pre-meiotic mitotic cell cycles or pre-meiotic S phase. Crossovers were completely abolished in *mei-41^29D^*; *mei-P22^103^* double mutants (*n* = 1754). One vial had two female progeny that were mutant for all markers on the *net–cn* chromosome except pr. These may have arisen from a double crossover in the adjacent *b–pr* and *pr–cn* regions, gene conversion of the *pr* mutation, or reversion of this mutation (an insertion of a *412* transposable element). Since these were in the same vial they likely represent a single pre-meiotic event. We conclude that the vast majority of crossovers observed in the *mei-41* null mutant females are meiotic in origin.

**Figure 1.**
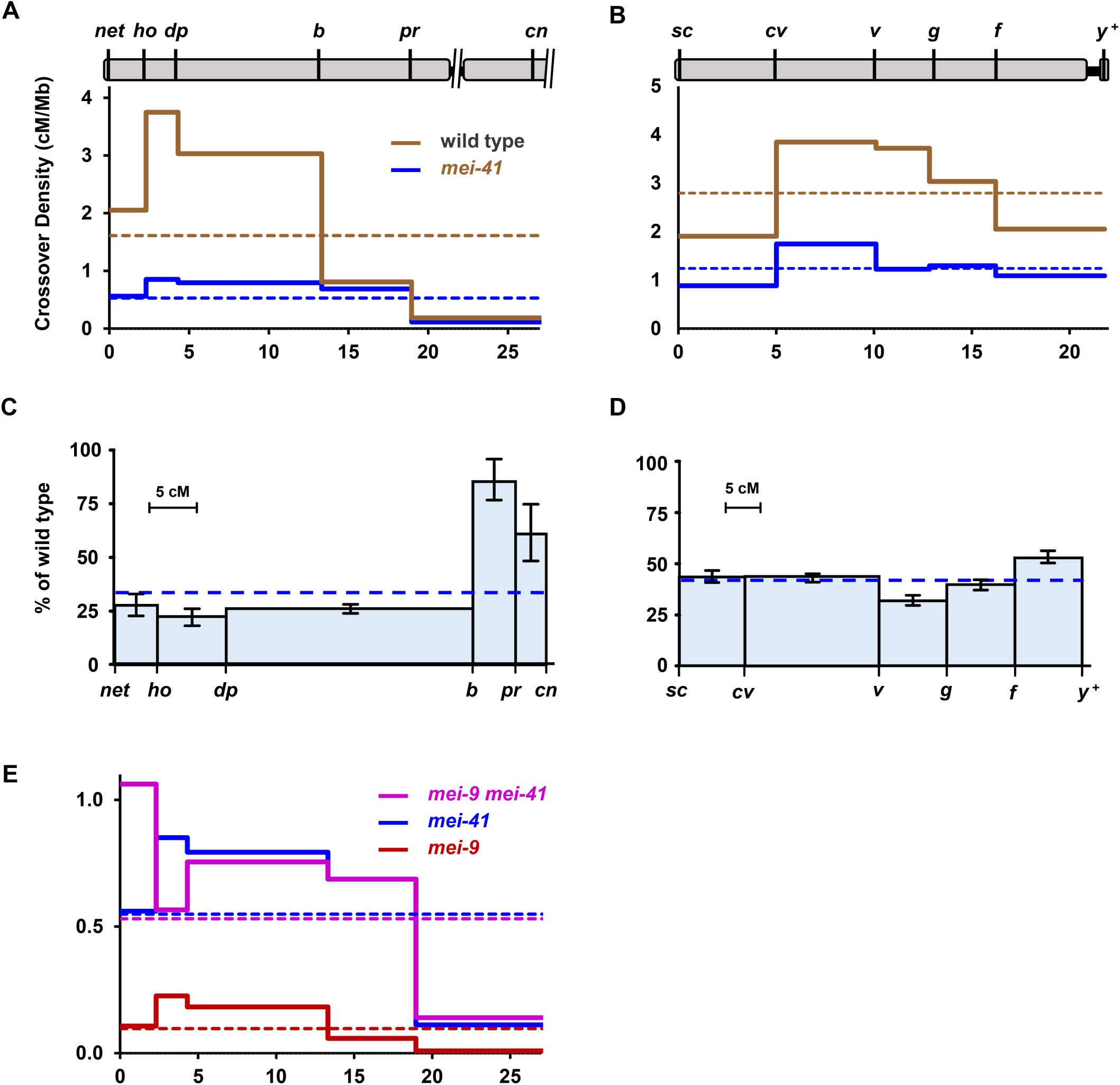
Reduction of crossing over in *mei-41* null mutants. (A) and (B) Crossover distribution on *2L* (A) and *X* (B) in *mei-41^29D^* mutants compared to wild type. Marker location indicated at top based on genome assembly position (Mb), excluding the centromere, unassembled pericentromeric satellite sequences, and transposable elements. Crossover density (solid lines) was determined for wild-type and *mei-41* mutant females. Dotted lines show mean crossover density across the entire region. (C) and (D) Crossing over on *2L* (C) and *X* (D) in *mei-41^29D^* mutants as a percentage of wild type. The X axis is scaled to genetic distance (cM) in wild-type females. Bars are 95% confidence intervals. (E) Crossover density in *mei-9* and *mei-41* single and double mutants. Note scale difference compared to (A). Wild type *2L*: *n* = 4222 progeny, 1943 crossovers; *mei-41 2L*: *n* = 7801 progeny, 1175 crossovers. Wild type *X*: *n* = 2179 progeny, 1367 crossovers; *mei-41 X*: *n* = 5174 progeny, 1396 crossovers; *mei-9*: *n* = 2433 progeny, 67 crossovers; *mei-9 mei-41*: *n* = 1059 progeny, 165 crossovers. Wild type and *mei 9* single mutant data are from Hatkevich *et al.* (2017), used with permission. Full datasets are in Tables S1 and S2. cM and cM/Mb, with 95% confidence intervals, are in Table S3.

Baker and Carpenter (1972) described crossover reduction in *mei-41* hypomorphic mutants as polar, with a more severe decrease in medial and distal regions of the chromosome than in proximal regions. This is also true in our null mutant: Although crossovers are significantly reduced in every interval, the average reduction in the three distal intervals is 75%, while in the two proximal intervals the decrease averages only 16% (Figure 1C). We also assayed crossing over across the entire *X* chromosome. Crossovers were reduced by an average of 57% on this chromosome; notably, the decrease was uniform across the entire chromosome, with no apparent polar effect (Figure 1B and 1D).

One hypothesis to explain the polar effect on recombination on *2L* is that there are region-specific requirements for Mei-41, with the protein being less important in proximal *2L*. We tested this hypothesis by assessing the dependence of crossovers on Mei-9, the catalytic subunit of the putative meiotic resolvase (Sekelsky *et al.* 1995). Meiotic crossovers are reduced by about 90% in *mei-9* mutants, suggesting that most or all crossovers generated in wild-type flies require Mei-9 (Figure 1E; Baker and Carpenter 1972; Sekelsky *et al.* 1995). However, in many mutants that affect meiotic recombination, including *Blm*, *mei-218*, and *rec*, crossovers are independent of Mei-9 (Sekelsky *et al.* 1995; Blanton *et al.* 2005; Hatkevich *et al.* 2017). Our interpretation is that when the meiotic crossover pathway is blocked because of loss of a critical component, repair is completed by alternative pathways that are independent of Mei-9 and other downstream meiotic recombination proteins. If Mei-41 is less important in proximal *2L*, then crossovers in these regions may remain dependent on Mei-9. We scored crossovers along *2L* in *mei-9^a^mei-41^29D^* double mutants (Figure 1E; we did not score the *X* chromosome because of the difficulty of recombining the *mei-9* and *mei-41* mutations and the *UASp::mei-41* transgene onto the multiple-marked chromosome, and because the requirement for Mei-41 appeared to be similar across the *X*). The total genetic map length was similar between *mei-9 mei-41* double mutants and *mei-41* single mutants (15.58 cM *vs.* 15.06 cM; *p* = 0.6679 by *χ*^2^ test comparing total crossovers and number of progeny scored) but significantly greater than that of *mei-9* mutants (2.8 cM; *p* < 0.0001). There was no apparent difference in requirement for Mei-9 between the proximal and distal intervals. We conclude that all crossovers generated in *mei-41* mutants are independent of Mei-9, regardless of chromosomal location. This suggests that loss of Mei-41 disrupts progression through the meiotic crossover pathway at all sites along the chromosome.

### The apparent polar effect on crossing over in mei-41 mutants can be explained by retention of the centromere effect

Compared to wild-type crossing over, the effects of loss of Mei-41 on meiotic crossing over is puzzling, as there seem to be substantially stronger effects in some regions of the genome than others, yet all crossovers in the mutant are independent of Mei-9. The conclusion that there is a polar effect on crossing over is based on comparing crossover frequencies in the mutant to those in wild-type females. Insight can also be gleaned by analyzing crossover distribution in the mutant in isolation. For example, Hatkevich *et al.* (2017) noted an apparently flat distribution of crossover in mutants lacking the Blm helicase. Their interpretation was that all crossover patterning is lost in *Blm* mutants, resulting in a distribution that reflects the DSB distribution.

Crossover distribution in *mei-41* mutants does not mimic that of *Blm* mutants, at least in proximal *2L*, suggesting that crossover patterning is not entirely lost in *mei-41* mutants. In wild-type flies, crossover density is substantially lower in the *pr–cn* interval than in any of the other nine intervals that we assayed (0.11 ±0.03 cM/Mb, versus 0.56 ±0.09 in the next lowest interval, *net–ho*). The *pr–cn* interval is noteworthy because it spans the centromere, so recombination is strongly influenced by the centromere effect. To determine whether this phenomenon is affected by loss of Mei-41, we calculated *CE* as a measure of the centromere effect (see Materials and Methods; Hatkevich *et al.* 2017). In wild-type females, if crossover density in the *pr-cn* interval were equal to the mean density across the entire region assayed, 643 crossovers would be expected; only 73 were observed (*p* < 0.0001), giving a *CE* value of 0.89. In *mei-41^29D^* mutants, 390 were expected but only 82 were observed, yielding a *CE* value of 0.79 (*p* < 0.0001). This high value of CE indicates that most or all of the centromere effect is intact in *mei-41^29D^* mutants, although the significant difference between *mei-41^29D^* and wild-type females (*p* = 0.0004) suggests that there may be mild amelioration in *mei-41* mutants.

The decrease in crossing over on the *X* chromosome does not appear to be polar (Figure 1C and 1D). The pericentric heterochromatin of the *X* chromosome spans about 19 Mb, compared to about 5 Mb on *2L* and 7 Mb on *2R*. This results in a much weaker centromere effect in the most proximal euchromatin of the *X* (Yamamoto and Miklos 1978). Thus, the lack of a polar decrease in crossing over on the *X* in *mei-41* null mutants may be because the entire region being analyzed is, with respect to distance from the centromere, equivalent to the distal half of *2L*.

The small chromosome *4* of *Drosophila melanogaster* never has meiotic crossovers in wild-type females (reviewed in Hartmann and Sekelsky 2017), but does have crossovers in *Blm* mutants (Hatkevich *et al.* 2017). Hatkevich *et. al.* argued that the absence of crossovers on *4* is due in large part to a strong centromere effect (about 4 Mb of heterochromatic satellite sequence between the centromere and the gene-containing region), and that it is loss of the centromere effect that permits crossing over on *4* in *Blm* mutants. We measured crossing over on *4* in *mei-41* mutants. We recovered no crossovers between markers at opposite ends of the gene-containing region of *4* (*n* = 5555; *p* < 0.0001 compared to *Blm*), consistent with our interpretation that the centromere effect is not lost in *mei-41* mutants.

### *Loss of crossover interference and assurance in* mei-41 *null mutants*

Given the apparent retention of the centromere effect on crossing over, we asked whether crossover patterning phenomena interference and assurance are impacted by loss of Mei-41. We calculated interference (*I*) using the method of Stevens (1936). Stevens defined *I* as 1-(*O*/*E*), where *O* is the number of double crossovers observed and *E* is the number of double crossover expected if the two intervals are independent of one another (see Materials and Methods). Thus, *I* = 1 indicates complete positive interference (no double crossovers observed) and *I* = 0 indicates no interference (the two intervals are independent of one another).

Values for *O*, *E*, and *I* are given in Figure 2A and 2B. On *2L*, the only pairs of adjacent intervals that have enough double crossovers to analyze interference are II/III (*ho–dp* and *dp–b*) and III/IV (*dp–b* and *b–pr*). In wild-type females, *I* was 0.93±0.05 between intervals II and III (*p* < 0.0001) and 0.64±0.15 between intervals III and IV (*p* = 0.0001). In *mei-41* mutants, we did not detect significant interference in the first pair of intervals (*I* = 0.26±0.52, *p* = 0.25; *p* < 0.0001), but interference appeared to be intact between the second pair of intervals (*p* = 0.9922 compared to wild type). The *dp–b* interval is typically not used in measuring interference because of its large size (>27 cM). Therefore, we reexamined interference within this region by subdividing it with another marker, *wg^Sp-1^*. Again, interference was strong in wild-type females but absent from *mei-41* mutants (Figure 2A, lower section). Using the same analysis of interference across the *X* chromosome, we found significant positive interference between every pair of adjacent intervals in wild-type females, but no detectable interference in *mei-41* mutants (Figure 2B).

**Figure 2.**
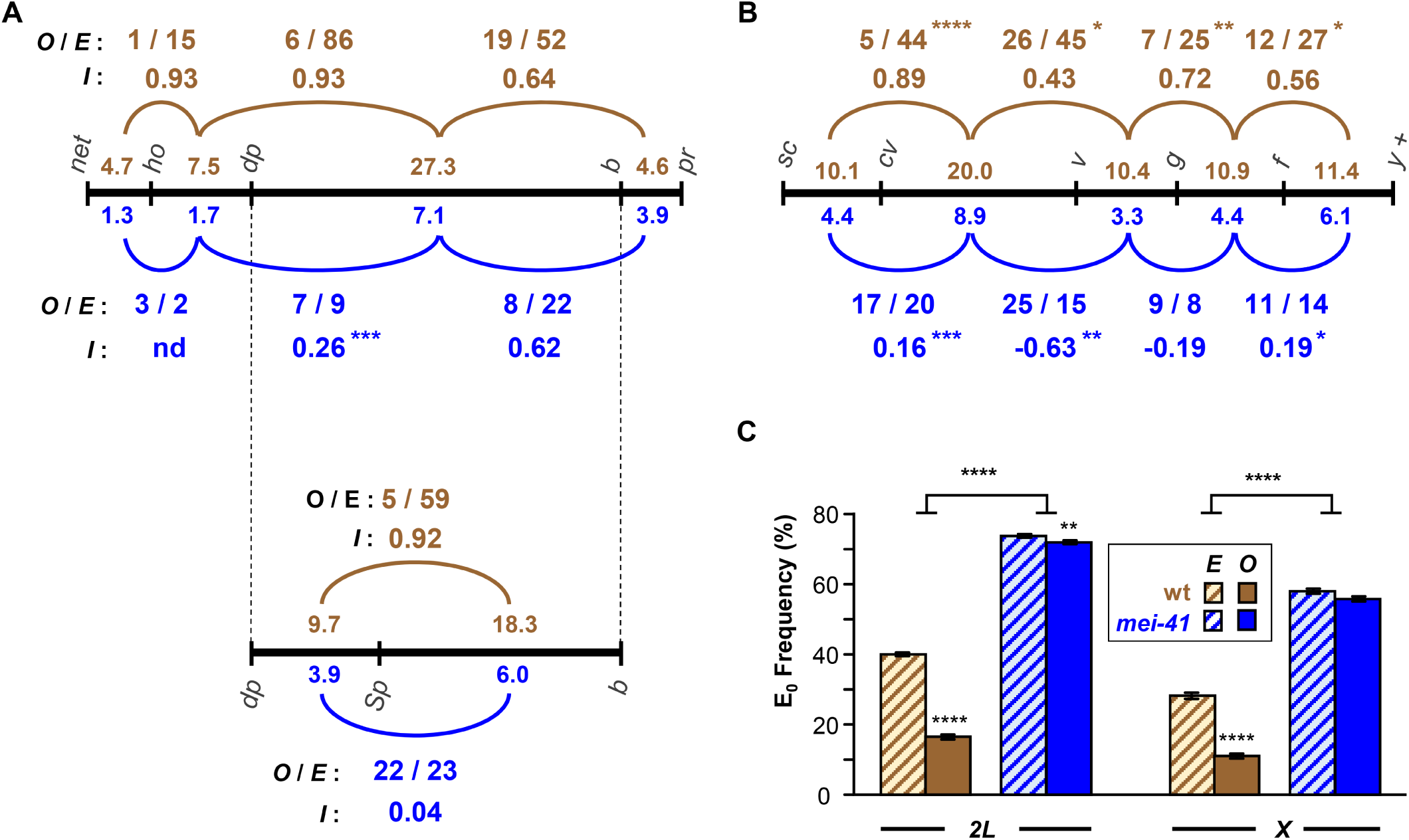
Interference and assurance in *mei-41* null mutants. (A) Interference on *2L*. Black line represents genetic map of markers used, with size of each interval (in cM) listed above the line for wild-type females and below the line for *mei-41* mutants. The *pr–cn* interval was omitted because it spans the centromere and because of low numbers of double crossovers in between this and the adjacent interval. Arcs represent pairs of adjacent intervals in which interference was tested. Above (for wild type, brown) or below (*mei-41*, blue) each arc is listed the number of observed double crossovers (*O*) and the number expected (*E*) if the two intervals are independent (no interference). Asterisks indicate *p* values for the difference between *O* and *E*. Stevens’ interference (*I*), which equals 1-(*O*/*E*), is also given. Asterisks on *I* values indicate *p* values for *χ*^2^ of *O* and *E* for wild type and mutant (see Materials and Methods). In a separate experiment, the large *dp–b* interval was further divided by the addition of *Sp* (*wg^Sp-1^*). (B) Similar analysis of interference on the *X* chromosome. The *f–y^+^* region spans the centromere, but since the marker on the right arm (*y^+^*) is hemizygous (*i.e.*, a duplication of the tip of *XL* onto *XR* on one ho-molog), all crossovers must be to the left of the centromere. (C) Crossover assurance assessed by comparing frequencies of E_0_ bivalents. Expected E_0_ frequency is based on Poisson distribution from the average number of crossovers per meiosis; observed frequencies were calculated using the method of Weinstein (1936). Statistical significance between expected and observed E_0_ frequencies determined via *χ*^2^ tests. Bars show 95% confidence intervals. Sample sizes for *2L* and *X* are given in Figure 1. For the *dp–Sp–b* experiment, *n* = 3325 flies, 928 crossovers for wild-type, 9740 flies, 972 crossovers for *mei-41*. * *p* < 0.05, ** *p* < 0.01, *** *p* < 0.001, **** *p*< 0.0001.

We used one additional method to assess the distribution of crossovers relative to one another. In many species, crossovers are distributed among bivalents such that the probability that any pair of homologous chromosomes does not receive a crossover is significantly lower than expected by chance, a phenomenon known as crossover assurance. It has been proposed that if there are sufficient well-spaced crossover-eligible intermediates, then coupling interference with a mechanism to achieve a specific number of crossovers per meiosis (within a narrow range) will produce crossover assurance (Zhang *et al.* 2014; Wang *et al.* 2015). In this model, assurance is merely an outcome of interference.

Extrapolating from our measurements of crossovers on *X* and *2L*, we estimate about two crossovers per meiosis in *mei-41* mutants. True assurance requires a minimum of three or five crossovers (one per major chromosome or arm, excluding 4); however, assurance among the residual crossovers could manifest as the two crossovers being on different chromosomes (or chromosome arms) more often than expected by chance. We compared the expected and observed frequency of meioses in which there were no crossovers (E_0_, for zero-exchange bivalent, frequency) on *X* or on *2L*. For expected E_0_ frequency we used the Poisson distribution expectation based on the average number of crossovers per meiosis. We used the method of Weinstein (1936) to transform counts of progeny that inherited parental, single crossover, double crossover, etc., chromatids to bivalent exchange classes (see Materials and Methods). In wild-type flies, the expected E_0_ frequency for the *X* chromosome is 0.285, but the observed frequency was 0.112 (Figure 2C). This demonstrates crossover assurance that is significant (*p* < 0.0001) but incomplete (11% of meioses have no crossovers between the X chromosomes), as has been observed in previous studies (*e.g.*, Koehler *et al.* 1996; Weinstein 1936). In *mei-41* mutants, reduced crossing over results in a higher expected E_0_ frequency (0.582), but unlike the case in wild-type flies, the observed frequency (0.572) was not significantly different (*p* = 0.3008). Similar results were obtained with the *2L* data (Figure 2C). For *2L*, the difference between observed and expected in *mei-41* mutants was significant (*p* = 0.0046), but given the small magnitude of the difference (0.740 expected, 0.720 observed), this may not be biologically meaningful.

Together, our data indicate that interference and assurance are significantly decreased or lost in *mei-41* mutants, though it is possible that crossovers in proximal *2L* retain interference.

## Discussion

We have demonstrated that the Gal4>*UASp* rescue successfully overcomes maternal-effect embryonic lethality of *mei-41* mutants, allowing us to perform meiotic crossover patterning analysis in null mutants. The crossover reduction in null mutants is more severe than that of the previously reported for hypomophic mutants (Baker and Carpenter 1972), but the non-uniform reduction in crossing over chromosome *2* is still present (Figure 1B).

We considered the hypothesis that the polar effect stems from differential requirement for Mei-41 in proximal and distal regions of the chromosome. However, in *mei-41* mutants, crossovers in all regions are independent of the presumptive resolvase Mei-9 (Figure 1C). Our interpretation is that this reveals an essential role for Mei-41 in carrying out meiotic recombination throughout the genome. In the absence of Mei-41 the meiotic pathway is disrupted and repair is completed by alternative pathways that neither require functions specific to the meiotic pathway nor result in properties normally associated with meiotic recombination, such as crossover patterning.

Since the apparent polar effect is observed on *2* but not on the *X*, we hypothesized that the centromere effect is retained in *mei-41* mutants. We calculated *CE*, a measure of how much crossover density in an interval deviates from the mean crossover density (Hatkevich *et al.* 2017), to compare the centromere effect between wild-type females and *mei-41* mutants. Although every interval deviates significantly from the mean in wild-type flies, the very strong deviation in the *pr–cn* region (*CE* = 0.89) is probably due primarily to the suppression of crossovers associated with proximity to the centromere. Direct confirmation of the presence of a centromere effect requires moving the sequences to be analyzed away from the centromere through chromosome rearrangement, a difficult experiment because of the need to have structural homozygosity combined with heterozygosity for markers. Nonetheless, our data suggest that a strong centromere effect is retained in the absence of Mei-41.

In contrast to the absence of a strong impact on the centromere effect, our analysis suggests that interference and assurance are significantly disrupted when Mei-41 is absent. It is notable that the only pair of intervals in which we detect significant interference includes the *b–pr* interval (IV), which is the closest interval to the centromere that we could analyze (in the two most proximal intervals, IV and V, there were too few double crossovers expected [three] and observed [two]). This could indicate that Mei-41 does have different functions in proximal regions than in other parts of the genome. However, given that there appears to be no interference within the adjacent interval III, the presence of interference between III and IV would require that crossover-eligible intermediates in III be subject to interference exerted by crossover designations in IV, while at the same time any crossovers designated within III not signal interference themselves. This seems unlikely, and perhaps indicates that there is some other effect or some idiosyncrasy associated with this particular interval.

Another argument that interference is reduced or absent in *mei-41* mutants is that the strength of interference is inversely proportional to genetic size of interval in which it is measured. Therefore, since genetic intervals become shorter in *mei-41* mutants, interference might be expected to become stronger. This expectation would not hold for recombination proteins required to generate crossovers after interference has occurred, such as the proteins that resolve crossover-designated intermediates into crossovers (the "crossover maturation" step in the models of Zhang *et al.* (2014)). Mei-9 and associated proteins are thought to be required for resolution (Baker and Carpenter 1972; Sekelsky *et al.* 1995; Yildiz *et al.* 2002). It is not meaningful to discuss interference in *mei-9* mutants, since the number of crossovers per meiosis is well below one (0.06), but the uniform discrease in crossovers across *2L* led Baker and Carpenter (1972) to conclude that Mei-41 acts earlier in crossover generation than Mei-9.

We believe the most parsimonious interpretation of our data is that loss of Mei-41 has little or no impact on the centromere effect but reduces or eliminates interference and assurance. We propose that crossover patterning in *Drosophila* occurs in a stepwise manner (Figure 3). Analysis of non-crossover gene conversion events mapped through whole-genome sequencing suggests that DSBs are, at a large scale, spread evenly throughout the assembled genome (Comeron *et al.* 2012; Miller *et al.* 2016; Hatkevich *et al.* 2017). The centromere effect is applied early by making some intermediates ineligible to enter the crossover pathway, with the probability of being affected in this way being related to distance to the centromere (Figure 3B). Subsequently, when any remaining crossover-eligible intermediate becomes crossover-designated, interference precludes nearby intermediates from also adopting this fate (Figure 3C).

**Figure 3.**
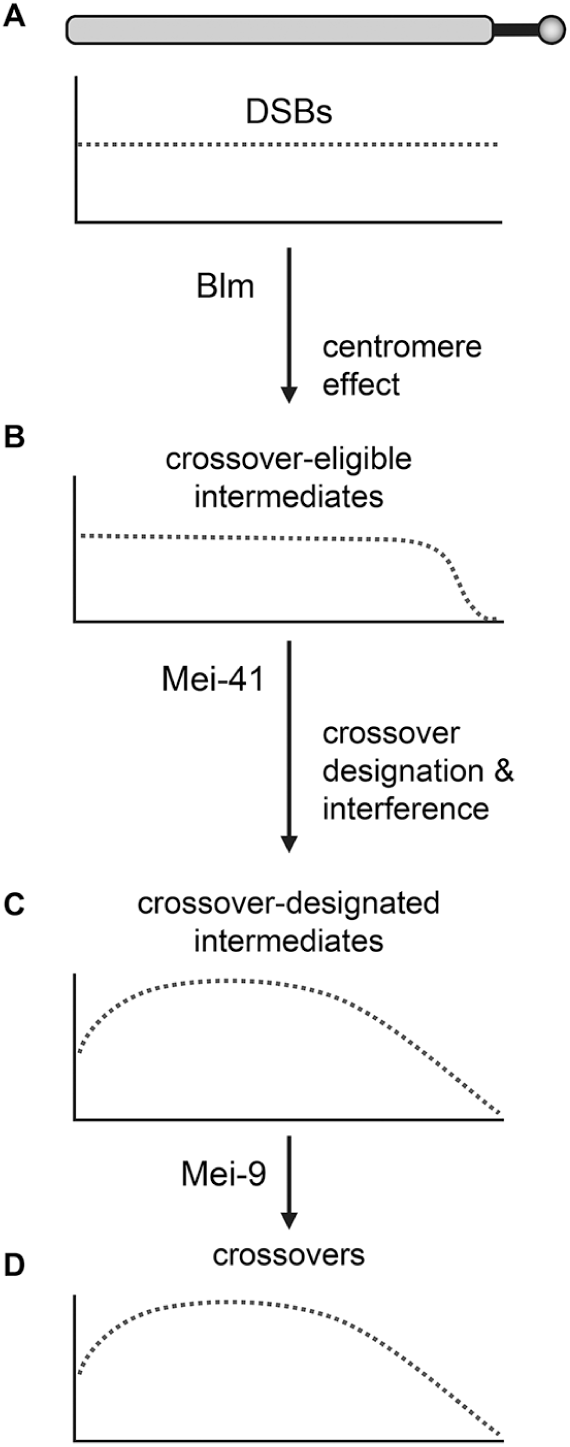
Model for progressive enforcement of crossover patterning. The drawing at the top represents a chromosome arm. Solid black line is pericentric satellite DNA, circle is centromere. (A) Based on whole-genome sequencing, the initial DSB distribution (dotted line) is flat at large scales (Comeron *et al.* 2012; Hatkevich *et al.* 2017; Miller *et al.* 2016); DSBs are excluded from the heterochromatic satellite DNA (Mehrotra and McKim 2006). The centromere effect revokes the eligibility of intermediates near the centromere from becoming crossovers. (B) This results in a distribution of crossover-eligible intermediates that is flat across much of the arm, then tailed as the centromere is approached. (The shape of the tailing is unknown; a sigmoidal drop is shown here for illustrative purposes.) Later, some intermediates are designated to become crossovers; these exert interference that discourages other intermediates over large distances from achieving crossover designation. The resultant distribution of crossover-designated intermediates (C) and crossovers (D) is approximately skew normal, with the degree of skew being proportional to the distance between the most proximal interval and the centromere. Blm, Mei-41, and Mei-9 are essential at different positions in the crossover pathway. Crossovers generated in mutants that lack these proteins are made outside the normal meiotic pathway, and are therefore Mei-9-independent. They are either unpatterned (*Blm* mutants, distribution similar to panel A), partially patterned (*mei-41* mutant, resembles panel B), or fully patterned (*mei-9* mutant, resembles panels C and D).

Given a uniform distribution of DSBs and the fact that each of the chromosome arms in *Drosophila* has about 1.0-1.3 crossovers per meiosis, interference alone will produce a crossover density that resembles a normal distribution (*e.g.*, the simulations in Zhang *et al.* 2014). The combination of a strong centromere effect and interference will yield a crossover density map that approximates a skew normal distribution (Figure 3D). Crossover distribution maps in *Drosophila* do resemble skew normal distributions, with much more skew on the major autosome arms than on the *X*, which also lacks a strong centromere effect (see Figure S2 in Comeron *et al.* 2012).

Blm helicase has been proposed to have a an essential function early in the meiotic recombination pathway (reviewed in Hatkevich and Sekelsky 2017). Loss of Blm results in an early exit from the meiotic pathway and completion of repair by alternative mechanisms. Since these alternative mechanisms do not involve patterning, the probability of becoming a crossover is the same for each intermediate, resulting in crossovers being evenly distributed across each chromosome arm. We propose that Mei-41 has some critical function after the centromere effect has been at least partially established. Loss of Mei-41 leads to exit from the meiotic pathway at this point. As with *Blm* mutants, every remaining intermediate has the same probability of becoming a crossover, to the crossover distribution in *mei-41* mutants is similar to the that in Figure 3B. Mei-9 is required only for maturation of crossover-designated intermediates into crossovers. Since this occurs after crossover designation, residual crossovers in a *mei-9* mutant are patterned like crossovers in wild-type flies, but there are far fewer crossovers because most intermediates that had been designated to become crossovers are instead processed into non-crossover products.

In many model organisms, a subset of crossovers do not participate in interference and are generated by a different resolvase than those that generate interfering crossovers (reviewed in Kohl and Sekelsky 2013). These “Class II” crossovers are sometimes defined as lacking interference or being unpatterned. In the model discussed above, this distinction is not always appropriate, at least in mutant situations. Rather, crossovers generated outside of the primary pathway may be unpatterned (as in *Blm* mutants), partially patterned (as in *mei-41* mutants), or patterned (as in *mei 9* mutants). The only features that these crossovers have in common is that they are generated outside of the normal meiotic crossover pathway, presumably through general DSB repair pathways that act to ensure there are no unrepaired DNA structures persisting until the meiotic divisions begin.

Our data provide little insight into the molecular function of Mei-41 in meiotic DSB repair pathway. In mitotic DSB repair, *mei-41* mutants have no observable defects in the early steps of homologous repair of DSBs I*e.g.*, resection, strand invasion, and repair synthesis), but Mei-41 appears to be required for the annealing and/or ligation steps of synthesis-dependent strand annealing (SDSA; LaRocque *et al.* 2007). Korda Holsclaw and Sekelsky (2017) hypothesized that Mei-41 activates Marcal1, which then catalyzes annealing of complementary sequences. SDSA promotes formation of non-crossover products, in contrast to the apparent role for Mei-41 in promoting crossovers during meiotic recombination. Nonetheless, Mei-41 might have a similar functions in mitotic and meiotic DSB repair if in the latter it activates a protein that catalyzes the annealing required for 2^nd^-end capture, a process that might occur after an early requirement for Blm helicase but prior to crossover designation (*e.g.,* models in Crown *et al.* 2014). Future studies to elucidate the role for Mei-41 might provide additional insights into meiotic crossover pathways and patterning.

## Acknowledgements

We thank Nicole Crown, Michaelyn Hartmann, and Talia Hatkevich for comments on the manuscript and Corbin Jones and Nadia Singh for advice on statistical analyses. This work was supported by a grant from the National Institute of General Medical Sciences (NIGMS) to JS under award 1R35GM118127.

---

**Table S1.**
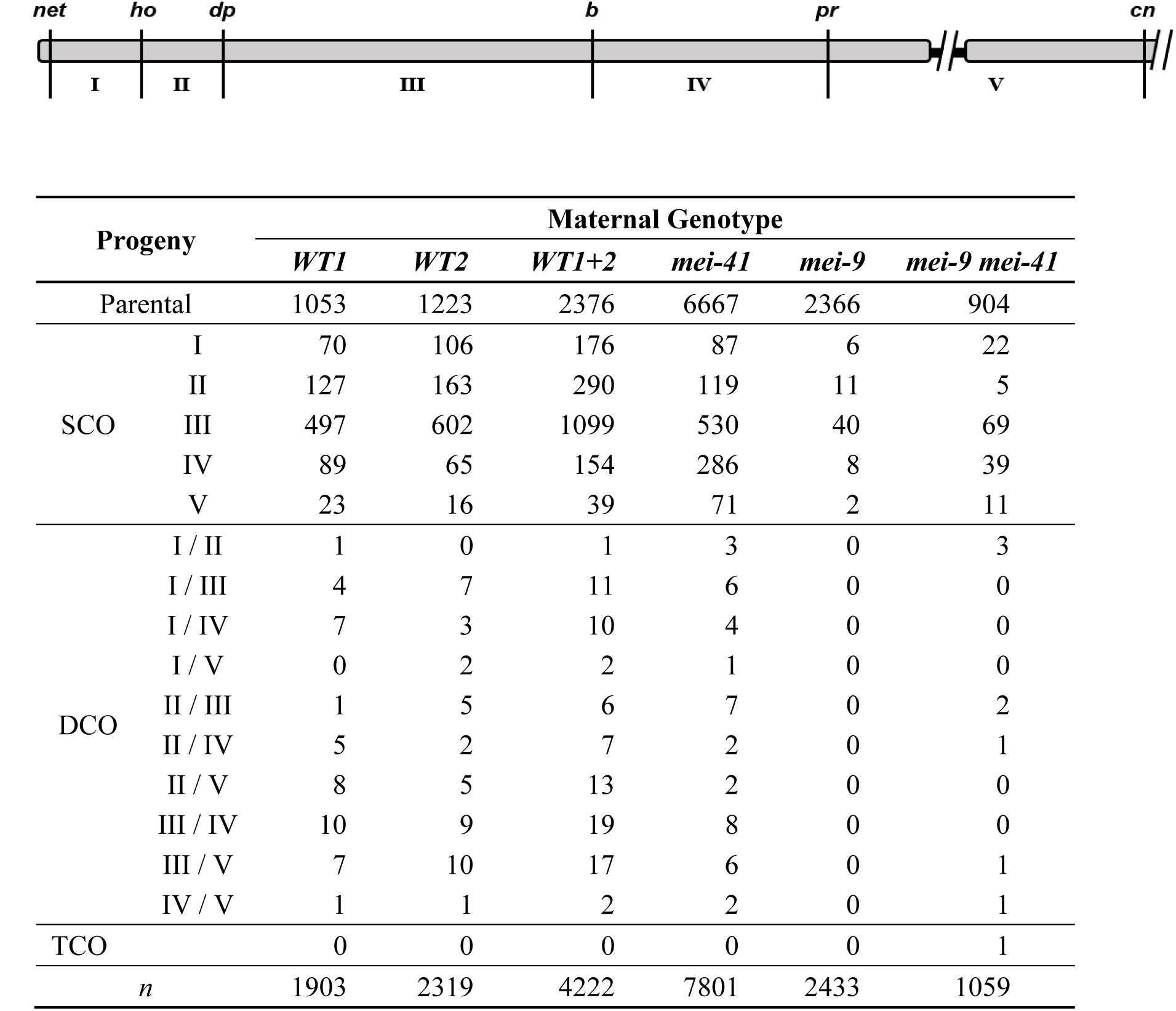
**Meiotic crossovers on chromosome *2L*.** Each row lists the number of total progeny from parental or single (SCO), double (DCO), or triple (TCO) crossover classes for wild-type (*WT*) and *mei-41* null mutants. Intervals I to V correspond to schematic above. Wild-type data were collected by different individuals in different years (see Material and Methods); the individual datasets and the summed set, which was used in all analyses, are given. The TCO in *mei-9 mei-41* was intervals I/II/IV. Wild-type data are from Hatkevich *et al.* (2017), used with permission (RightsLink license 4217090536151, 27 Oct 2017).

**Table S2.**
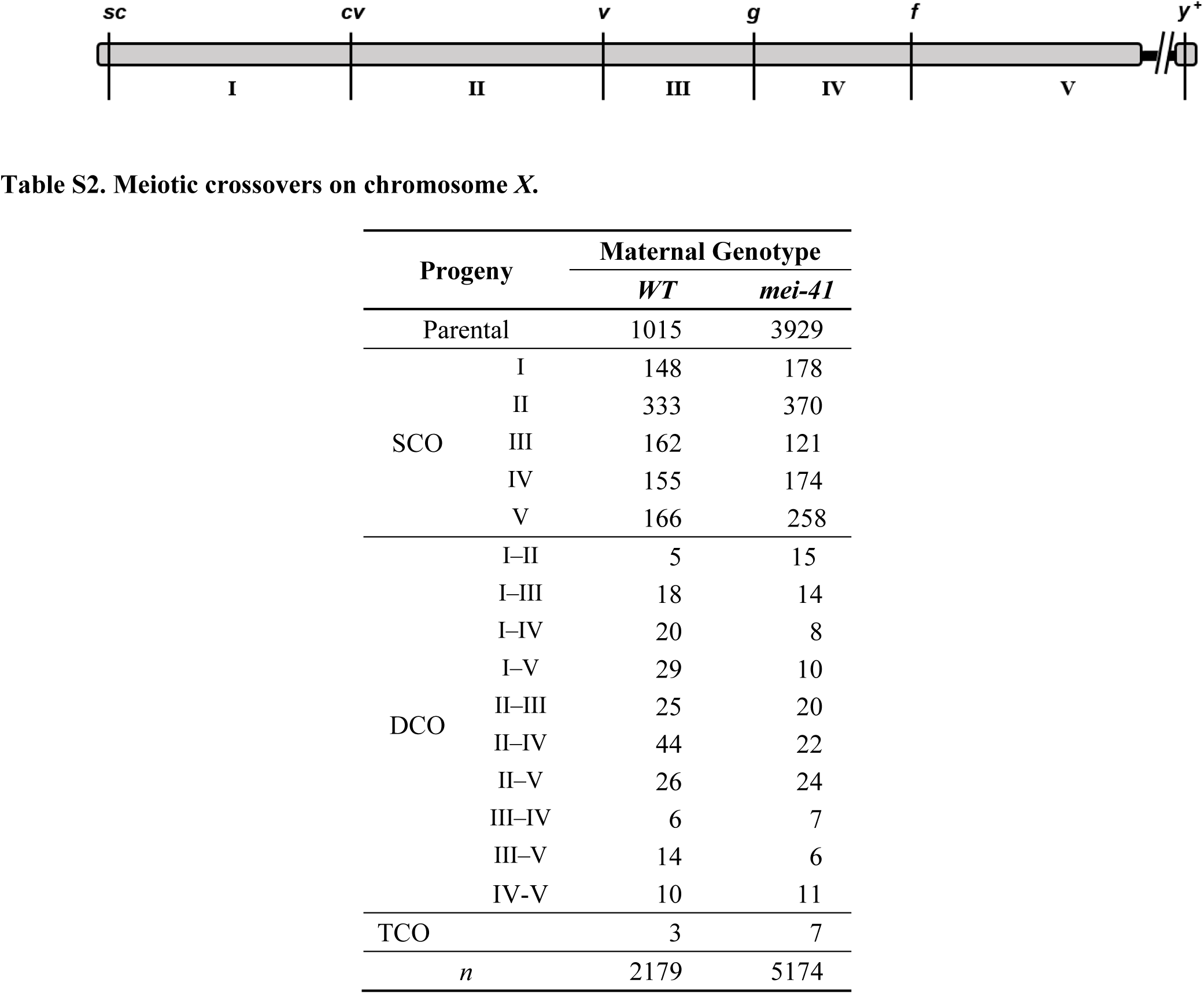
**Meiotic crossovers on chromosome *X*.** Each row lists the number of progeny from parental or single (SCO), double (DCO), or triple (TCO) crossover classes for wild-type (*WT*) and *mei–41* null mutants. Intervals I to V correspond to schematic above. The TCOs in wild/type flies were one each in intervals (II/III/V), (II/IV/V), and (III/IV/V). TCOs in *mei/41* were one each intervals (I/II/III), (I/II/IV), (I/II/V), (I/III/V), and (II/IV/V) and two each in intervals (II/III/IV) and (II/IV/V). Wild/type data are from Hatkevich *et al.* (2017), used with permission (RightsLink license 4217090536151, 27 Oct 2017).

**Table S3. Genetic distances and crossover densities.**The top section of each table gives calculated genetic distances (in cM, with 95% confidence intervals (CI); see Materials and Methods) for the five intervals on *2L* (see Tables S1) and *X* (see Table S3). The rightmost column has the summed distance across all five intervals. The lower two sections give crossover density (cM/Mb) calculated without including transposable elements (middle) or including transposable elements (bottom). Transposable element lengths are from the *Drosophila melanogaster* reference genome and are not necessarily the same in the chromosomes we used.

**Table S3a.**
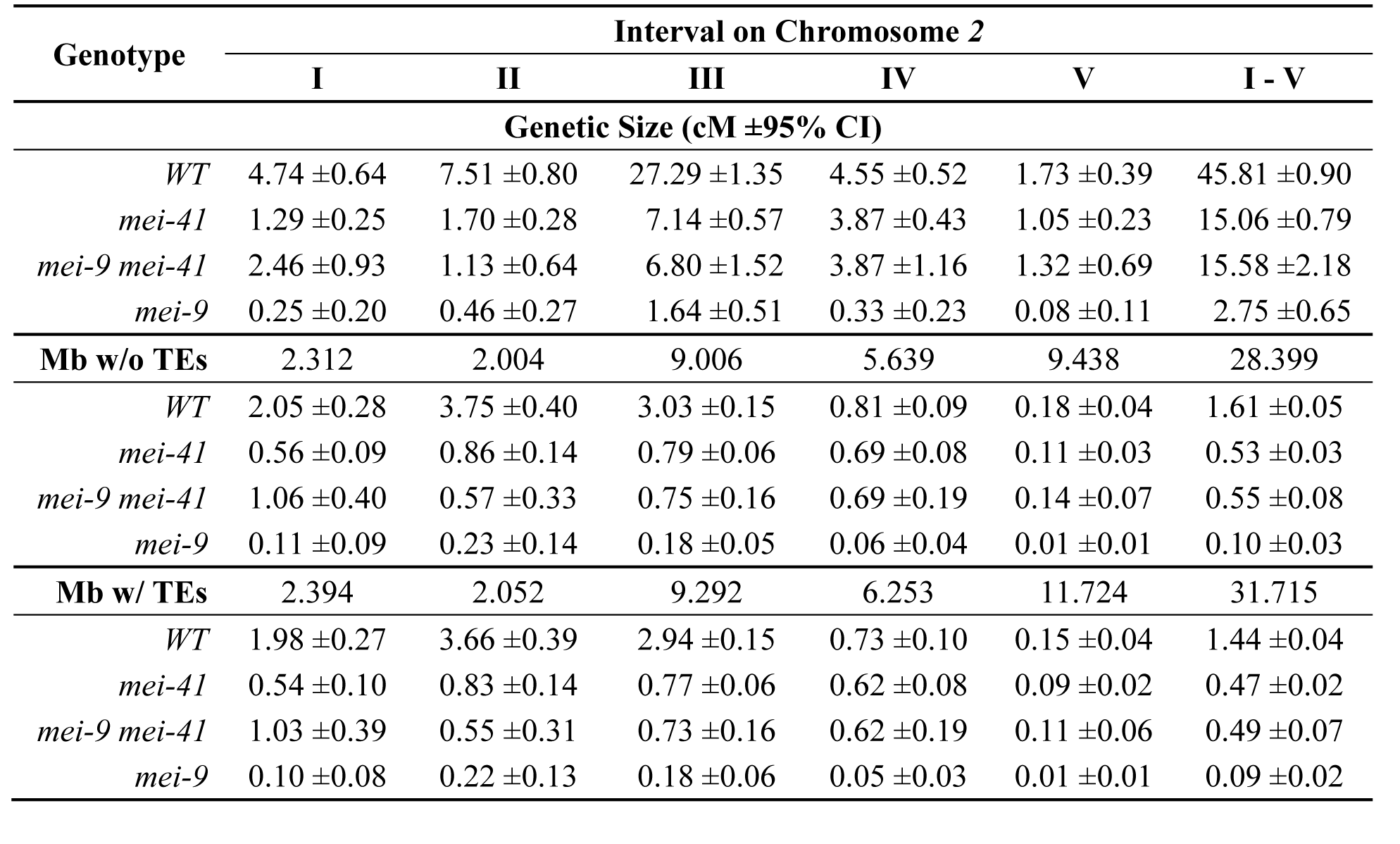
Genetic distances and crossover densities on chromosome *2*.

**Table S3b.**
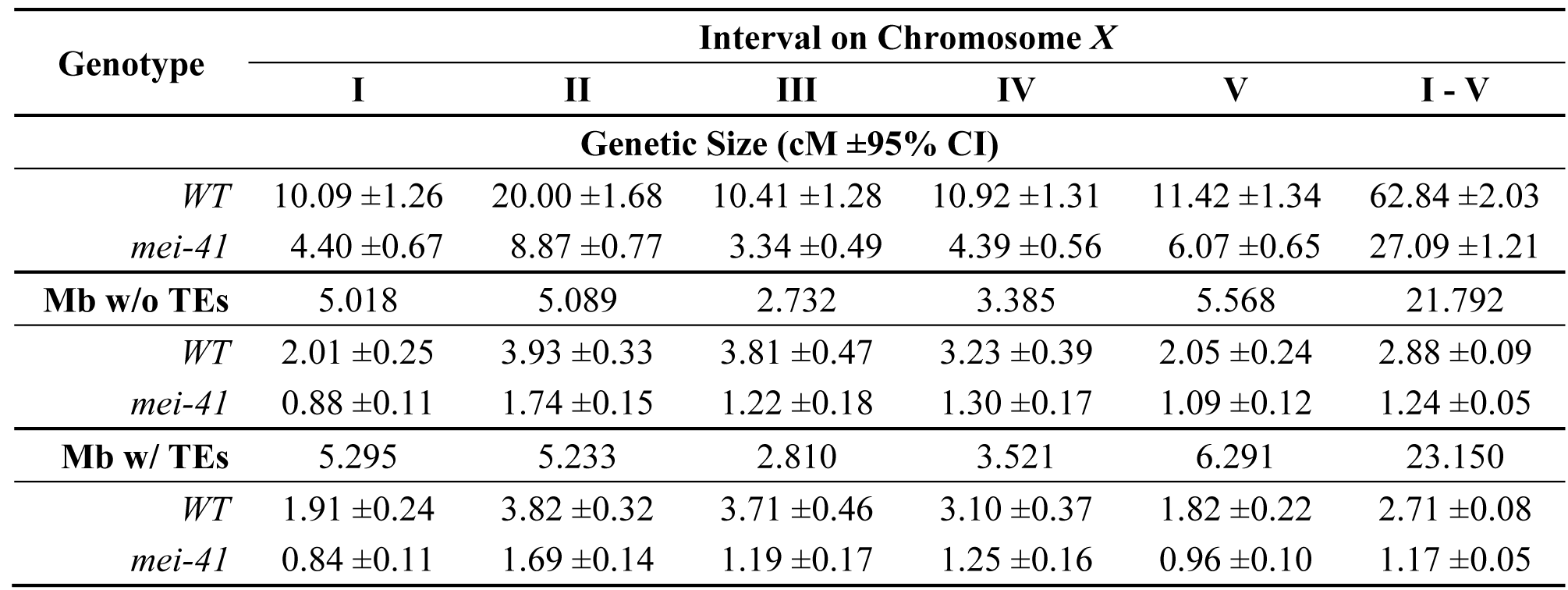
Genetic distances and crossover densities on chromosome *X*

**Table S4.**
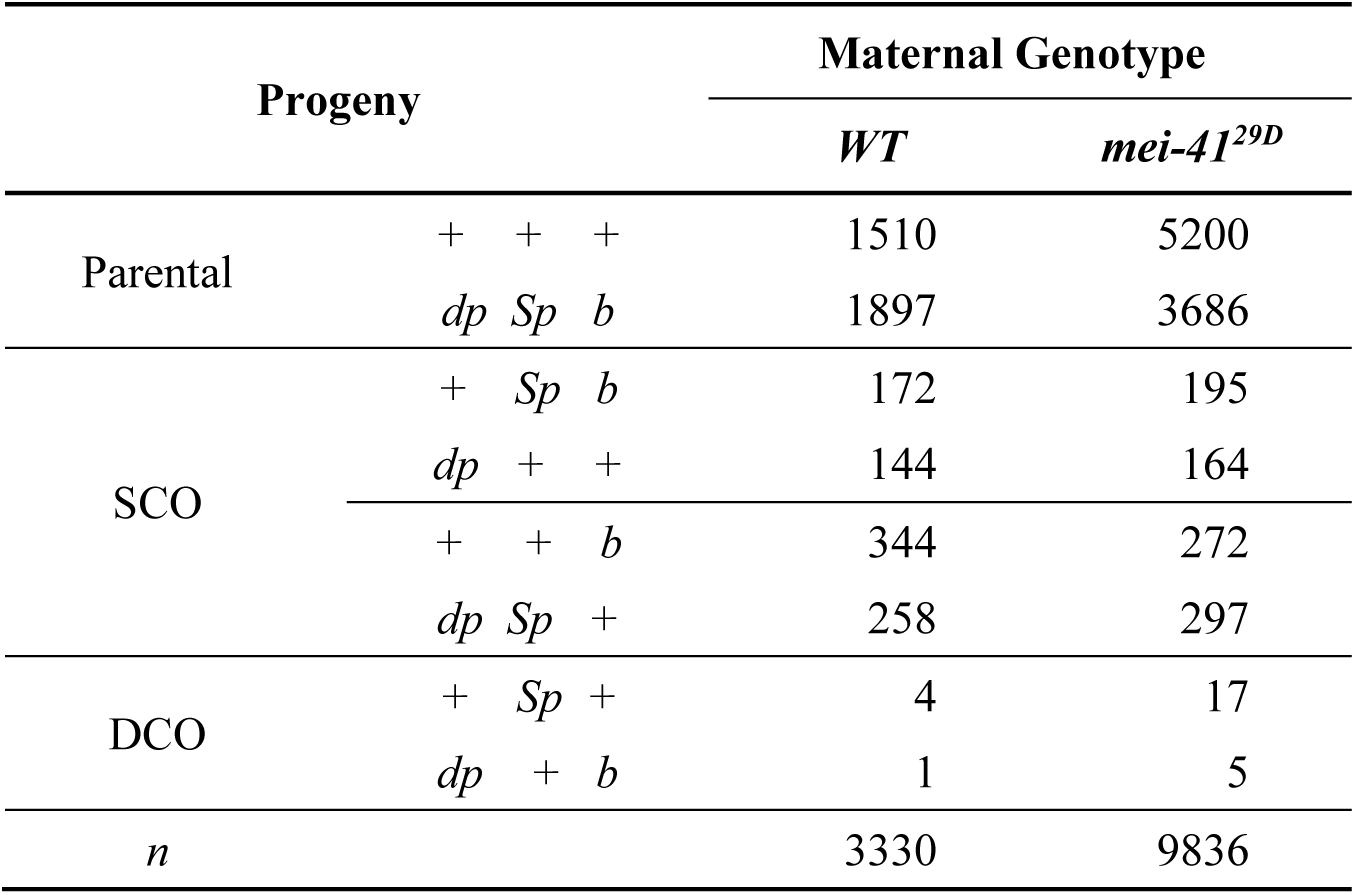
**Table S4. Progeny counts from *dp–Sp–b* interference experiment.** Each row lists the number of progeny from parental and single (SCO), or double (DCO) classes for wild-type (*WT*) and *mei-41* null mutants.

**Table S5a.**
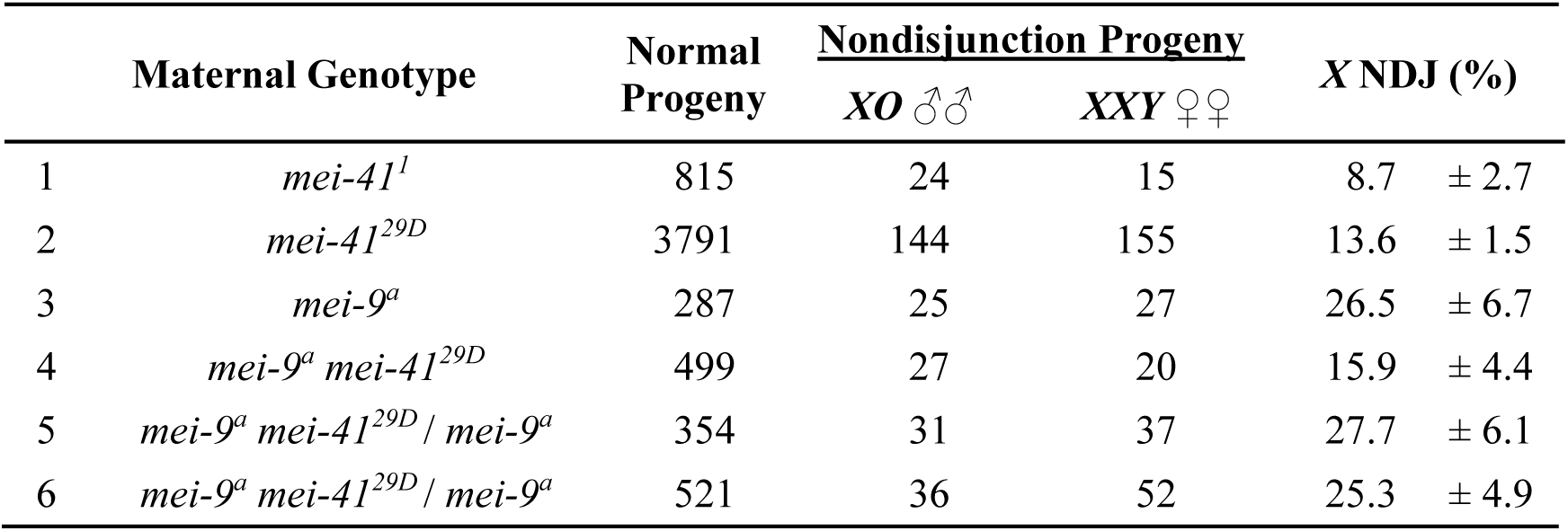
***X* nondisjunction.** *X* chromosome nondisjunction (NDJ) was scored as described in Materials and Methods. Genotypes 1-4 were homozygous for the indicated mutant alleles. All *mei-41^29D^* experiments had the *M*{*UASp::mei-41*} and *P*{*matα::GAL4*} transgenes described in the text. Genotypes 5 and 6 were made by crossing each of the two stocks that were used to generate *mei-9 mei-41* double mutants to *mei-9*^*a*^ single mutants to test for the presence of *mei-9*^*a*^ in the stock. The males used to generate genotype 5 were *y M*{*UASp::mei-41*} *mei-9^a^ mei-41^29D^* on the *X* chromosome. The males used to generate genotype 6 were *y mei-9^a^ mei-41^29D^* on the *X*. Statistical analyses are in Table S5b, below. *p* values are not corrected for multiple comparisons, but such corrections would not change any conclusions.

**Table S5b.**
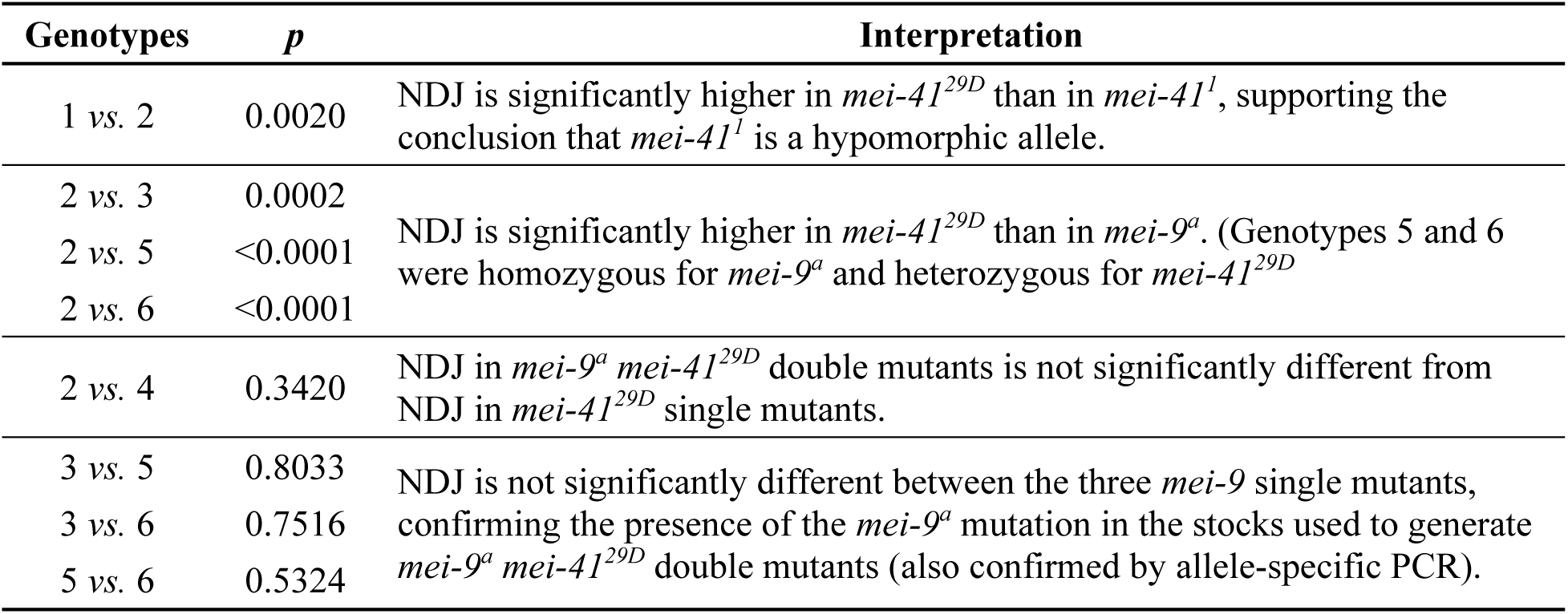
**Statistical comparisons of *X* nondisjunction.** Methods of Zeng *et al.* (2010) was used to calculate *X* nondisjunction (NDJ) with 95% confidence intervals and to calculate *p* values based *Z* tests. *p* values are not corrected for multiple comparisons, but such corrections would not change any conclusions.

